# QuadST: A Powerful and Robust Approach for Identifying Cell–Cell Interaction-Changed Genes on Spatially Resolved Transcriptomics

**DOI:** 10.1101/2023.12.04.570019

**Authors:** Jinmyung Choi, Michelle E. Ehrlich, Panos Roussos, Pei Wang, Guo-Cheng Yuan, Xiaoyu Song

## Abstract

Spatially resolved transcriptomics (SRT) have enabled profiling spatial organization of cells and their transcriptome in situ. Various analytical methods have been developed to uncover cell-cell interaction processes using SRT data. To improve upon existing efforts, we developed a novel statistical framework called QuadST for the robust and powerful identification of interaction-changed genes (ICGs) for cell-type-pair specific interactions on a single-cell SRT dataset. QuadST is motivated by the idea that in the presence of cell-cell interaction, gene expression level can vary with cell-cell distance between cell type pairs, which can be particularly pronounced within and in the vicinity of cell-cell interaction distance. Specifically, QuadST infers ICGs in a specific cell type pair’s interaction based on a quantile regression model, which allows us to assess the strength of distance-expression association across entire distance quantiles conditioned on gene expression level. To identify ICGs, QuadST performs a hypothesis testing with an empirically estimated FDR, whose upper bound is determined by the ratio of cumulative associations at symmetrically smaller and larger distance quantiles simultaneously across all genes. Simulation studies illustrate that QuadST provides consistent FDR control and better power performance than other compared methods. Its application on SRT datasets profiled from mouse brains demonstrates that QuadST can identify ICGs presumed to play a role in specific cell type pair interactions (e.g., synaptic pathway genes among excitatory neuron cell interactions). These results suggest that QuadST can be a useful tool to discover genes and regulatory processes involved in specific cell type pair interactions.

## Introduction

Cell-cell interactions underlie the organization of cells and can affect the function of tissues that are made up of diverse cell types ^1^. Recent advances in single-cell sequencing technologies have enabled the study of diverse cell types and their interactions^2^. In particular, spatially resolved transcriptomics (SRT) has gained great interest^1^ because it provides transcriptomic profiles of cells along with their spatial organization in native tissues. Such spatial information is vital to discover cell-cell interactions underlying development, homeostasis, and disease since cell-cell interactions can occur by signaling processes (e.g., juxtacrine and paracrine signaling) at a short distance (0 to 200 *μm*)^3^. The identification of cell-cell interactions pertinent to a specific context requires the development of analytical methods in conjunction with advances in SRT technology.

There exist various efforts in developing computational and statistical methods based on SRT data for the identification of cell-cell interactions and genes involved in cellular interaction processes ^4–10^. Giotto^4^ and NCEM ^5^ allow the identification of interaction-changed genes (ICGs) in a specific cell type pair’s interaction. SVCA ^6^ and MISTy ^7^ focus on identifying ICGs without specifying a particular cell type pair, and SpatialDM^8^, stLearn^9^, and SpaOTsc^10^ identify the patterns of cell-cell interactions using ligand and receptor pairs or a set of highly variable genes. We found that these statistical methods have a few common drawbacks that can be further improved (Supp. Table 1): i) Putative cell-cell interaction partners are predetermined by a particular distance threshold of the model, which can be inaccurate and result in reduced power. ii) Excessive zeros arising from the single-cell transcriptome are not considered in modeling. iii) Covariates that may introduce confounding effects in the associations between gene expression level and cell-cell distance cannot be explicitly specified in the modeling process. iv) The inference procedure lacks appropriate false discovery rate (FDR) control in the presence of correlation and confounders. v) Inference relies on known gene functions, such as ligands and receptors.

As an effort to tackle these problems, we propose a novel statistical framework for the identification of genes involved in cell-type-pair-specific interactions. The framework consists of the following features: i) modeling cell-cell distance as a continuous response variable, ii) including information about excess zeros in the cell-cell interaction model, iii) applying an empirical FDR control procedure applicable in the presence of correlation and confounding variables in transcriptome data, iv) using a regression framework to naturally handle covariate adjustment, and v) being unbiased regarding known gene function, cell type pair, and cell-cell interaction distance. These features contribute to the robust and powerful identification of genes involved in specific cell type pair interactions.

## Results

We developed a robust and powerful statistical framework called QuadST to identify interaction-changed genes (ICGs) for cell-type-pair specific interactions on single-cell SRT data. QuadST’s framework consists of 3 key steps (Fig. 1a and Methods). First, we define cell types, pair cell types of interest, and calculate the cell-cell distance of each cell type pair from SRT data. Each cell type pair consists of what we call “anchor” and “neighbor” cells, and cell-cell distance refers to the distance from each anchor cell to the closest neighboring cell. Second, we used a quantile regression model to test the association between cell-cell distance and gene expression level at a grid of evenly distributed cell-cell distance quantile levels. Additional variations of gene expression from technical and biological sources other than cell types can be adjusted as covariates. Third, we identified ICGs for a given anchor-neighbor cell pair based on hypothesis testing with an empirically estimated FDR, whose upper bound is determined by the ratio of cumulative associations at symmetrically smaller and larger distance quantiles simultaneously across all genes. This provides robust and efficient FDR control in the presence of correlations (e.g., gene-gene correlations) and confounders in the transcriptome data. Note that ICGs are genes within anchor cells showing a significant change in expression level with respect to anchor-neighbor cell pair distance (Methods).

**Figure 1.**
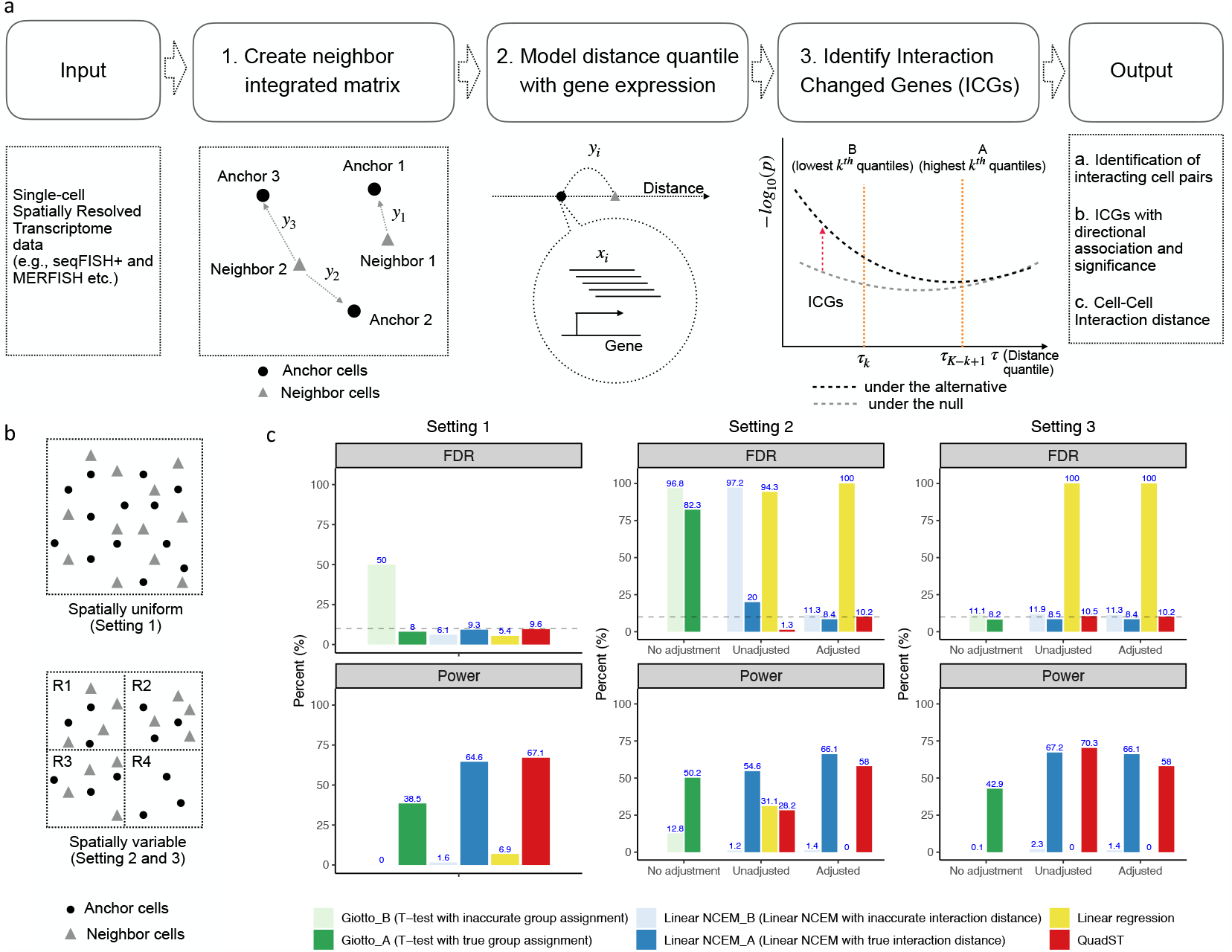
Overview of QuadST and its performance on simulated datasets. a. The statistical framework of QuadST consists of 3 key steps: (1) create an anchor-neighbor integrated matrix that includes anchor-neighbor cell-cell distance, anchor cell gene expression levels, and covariates, (2) model anchor-neighbor cell-cell distance quantile with the gene expression level of anchor cells, and (3) identify ICGs for a given anchor-neighbor cell pair based on a hypothesis testing with empirical distribution applicable in the presence of correlation and confounding variables. b. Simulation settings: The top panel illustrates Setting 1 with no spatial variation in cell composition and gene expression level. The bottom panel illustrates Setting 2 with spatial variation in cell composition and expression level and Setting 3 with spatial variation only in cell composition. c. Comparison of the power and FDR between QuadST and other methods on three different simulation settings. Across all settings, we simulated a ground truth set of interacting cells and ICGs from an anchor-neighbor cell pair. For each setting, we show the FDR and power averaged over 10 repeated simulation instances for each method. For Settings 2 and 3, we include the FDR and power calculated after adjusting for the spatial region (R4 in the bottom panel of b).

We designed simulation studies to evaluate the power and FDR of QuadST in comparison with other methods. We considered three settings representing various spatial organization of cells and their expression levels (Fig. 1b and Methods). The settings represent various spatial organization of cells and their expression levels. In Setting 1, there is no spatial variation in cell composition and gene expression level. In Setting 2, there is spatial variation in both cell composition and gene expression level. Such spatial variation can influence both cell-cell distance and gene expression level. As a result, the spatial region can serve as a confounder in distance-expression association. In Setting 3, there is a spatial variation only in cell composition. The spatial variation in cell composition can influence cell-cell distance without affecting gene expression levels. As such, the spatial region does not serve as a confounder. Across all settings, we first determined interacting cells based on proximity between two cells of different cell types and then introduced expression changes to 10% of genes in the interacting anchor cells. The particular choice of parameters used to simulate the ground truth, such as a certain threshold of cell proximity and a fraction of genes involved in cell-cell interactions, does not affect the generality of our simulation results.

We compared the performance of QuadST with that of three other methods. The first two methods, Giotto ^4^ and NCEM ^5^, are chosen from existing methods (Supplementary Table S1). This is because Giotto and NCEM allow the identification of ICGs for a particular cell type pair’s interaction so that the results are directly comparable to those of QuadST. The third method is replacing the quantile regression with the linear regression in the QuadST pipeline called “Linear regression”. Like QuadST, this method models cell-cell distance as a response and gene expression level as a predictor. However, it models mean distance as a response, whereas QuadST models entire distance quantiles as a response. Thus, it can provide a comparison between modeling entire distance quantiles and modeling mean distance only as a response variable. Note that both Giotto and NCEM require a predetermination of adjacency threshold to divide cells into a dichotomous variable of adjacent vs. nonadjacent cells for putative interactions between two cells, which differs from that of QuadST. We considered two different ways of choosing the adjacency threshold for both Giotto and NCEM. In each case, the distance threshold corresponds to either i) matching the ground truth interaction distance or ii) deviating from the ground truth interaction distance (Methods). In the comparison of the FDR and power, we included the FDR and power calculation with and without adjustment of the spatial variation in Settings 2 and 3. NCEM, Linear regression, and QuadST allow us to model such spatial variation as a covariate and adjust for it in the data analysis, whereas Giotto allows us to model and adjust it in the data preprocessing step.

Fig. 1c shows the FDR and power calculation for each of the compared methods across three different simulation settings. The following are the key observations. First, QuadST consistently controls FDR at a prescribed level (10%) (Fig. 1c top row), even in the presence of an unadjusted confounder. On the other hand, other methods can show inflated FDR: i) Linear regression suffers from a highly inflated FDR in Settings 2 and 3, regardless of covariate adjustment (Fig. 1c top row; middle and right panels). This occurred due to the presence of a single peculiar false-positive ICG, which remained significant even after covariate adjustment in both settings. ii) Giotto and NCEM can control the FDR at a prescribed level only after covariate adjustment, indicating their lack of FDR controlling capacities under unadjusted confounders. This finding suggests that QuadST is the only method that avoids detecting false positives in the presence of an unadjusted confounder. Second, QuadST achieves similar or better power compared to other methods (Fig. 1c bottom row). On the other hand, other methods can show poor or variable power: i) The power of Linear regression remains consistently poorer than that of the other methods in all settings (Fig. 1c bottom row). ii) The power of Giotto and NCEM decreases as the distance threshold used in the model deviates from the ground truth interaction distance, comparing Giotto *A* vs. Giotto *B* and NCEM *A* vs. NCEM *B* regardless of the settings (Fig. 1c bottom row). It is worth noting that in Setting 3 (Fig. 1c bottom row; right panel), where the spatial region is not a confounder, overadjustment of this variable does not largely affect the power of QuadST. In summary, we illustrated that QuadST consistently provides robust FDR control and high study power across various simulation settings.

To demonstrate the applicability of QuadST, we analyzed two SRT datasets^11,12^ based on seqFISH+ (sequential Fluorescence In Situ Hybridization plus) and MERFISH (Multiplexed Error Robust Fluorescence In Situ Hybridization) technologies. Both seqFISH+ and MERFISH datasets provide spatial organizations of cells and their gene expression profiles obtained from imaging postnatal mouse brain regions without known abnormality. The seqFISH+ dataset profiled a transcriptome-wide geneset of 10,000 genes obtained in hundreds of cells, while the MERFISH dataset profiled a curated geneset of about 400 genes in the order of 10,000 cells.

After data preprocessing (Methods), we focused on analyzing the following subsets from each dataset: the seqFISH+ dataset consists of gene expression levels of 1,235-2,500 genes in each of 511 cells from 23-day old male mouse (C57BL/6J) cortex. The MERFISH dataset consists of gene expression levels of 280 genes in each of 13,745 cells from 4-week-old female mouse (C57BL/6J) cortical layers II-V. The cells in each dataset were categorized into 6 and 8 major neuronal and nonneuronal cell types by original studies, respectively. We used a uniform manifold approximation and projection (UMAP) plot (Supp. Fig. S1a) to examine the effect of covariates on the gene expression profiles of cells from each dataset: cell types and field of view (FOV) from the seqFISH+ dataset and cell types and cortical layers from the MERFISH dataset. We found that cells cluster primarily by cell type in both datasets and secondarily by FOV and cortical layers in seqFISH+ and MERFISH datasets respectively, which were adjusted for in our analysis.

We ran QuadST for a total of 36 and 64 pairs of the same and different cell types in each of the seqFISH+ and MERFISH datasets, respectively. We identified a total of 639 and 149 unique ICGs across 69.4% (25 out of 36) and 68.8% (44 out of 64) of anchor-neighbor cell pairs in the seqFISH+ and MERFISH datasets, respectively (Fig. 2a). The detected ICGs show both positive and negative directions of association, indicating that they can either increase or decrease expression levels with cell-cell distance (Fig. 2b). We note that 96% (271/281) and 100% (16/16) of ICGs found in excitatory neuron-excitatory neuron cell interactions were positively associated in both the seqFISH+ and MERFISH datasets.

**Figure 2.**
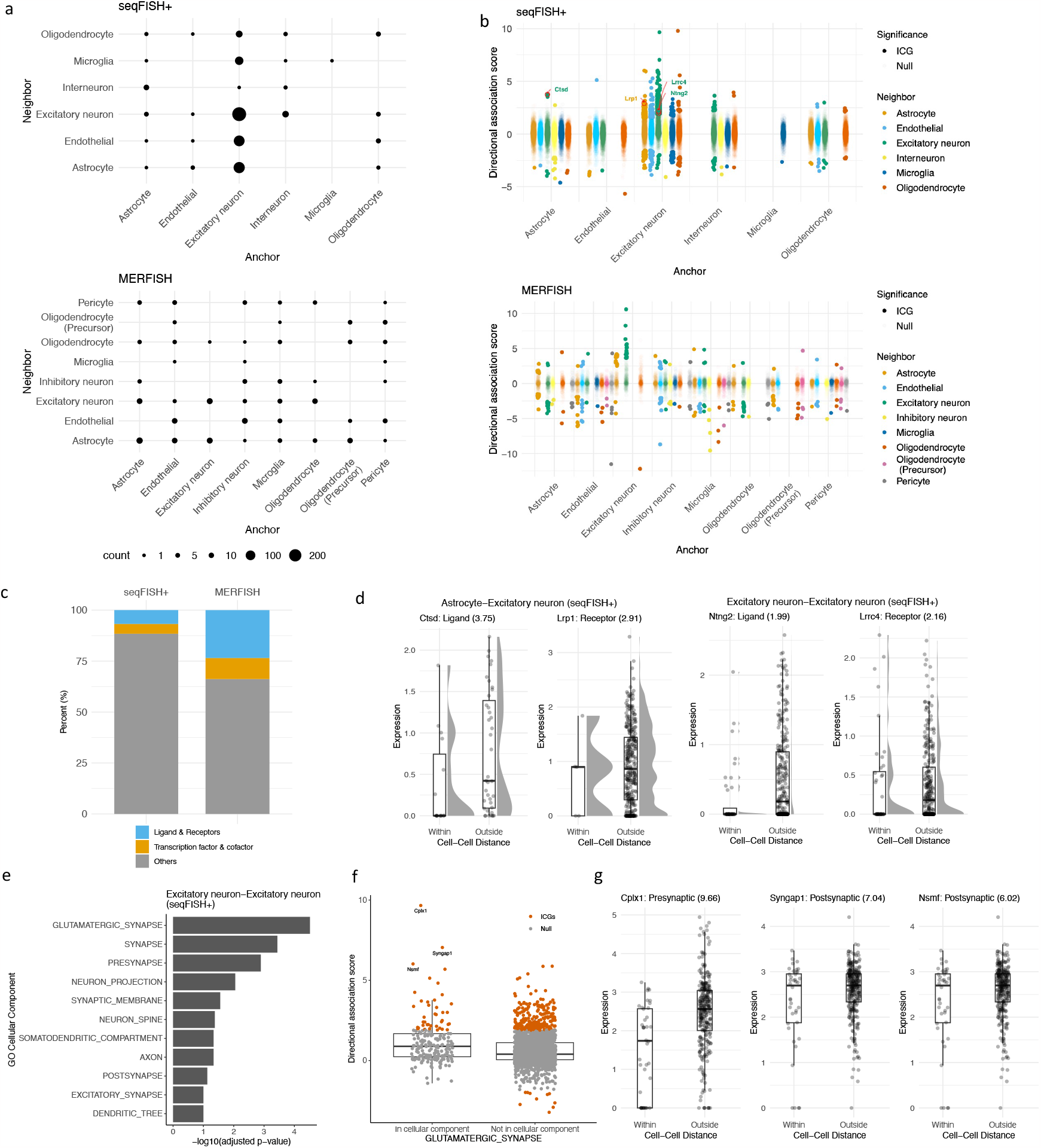
Application of QuadST on two SRT datasets obtained from mouse brains. a. Number of ICGs among all anchor-neighbor cell pairs. We identified a number of ICGs among major neuronal and nonneuronal cell type pairs in both SRT datasets. b. Directional association scores of ICGs and all other genes among all anchor-neighbor cell pairs. The ICGs show both positive and negative directions of association, indicating that their expression levels either increase or decrease with cell-cell distance. Highlighted are two ligand–receptor ICG pairs, *Ctsd* -*Lrp1* and *Ntng2* -*Lrrc4*, in the seqFISH+ dataset. *Ctsd* (ligand) and *Lrp1* (receptor) were detected as ICGs in astrocyte-excitatory neurons and excitatory neuron-astrocyte cell pairs, respectively. *Ntng2* and *Lrrc4* are detected ICGs in the excitatory neuron-excitatory neuron cell pair. c. Percentage of functional categories of ICGs detected in seqFISH+ and MERFISH. d. Distance-expression profiles of highlighted ligand-receptor pairs are shown in (b). We show the boxplot along with a half-eye plot (showing the density of jittered points) of the gene expression levels of each ligand and receptor within and outside of the cell-cell interaction distance. Shown in parentheses are the directional association scores. e. GO cellular component enrichment analysis of ICGs detected in excitatory neuron-excitatory neuron cell pair interactions. We found that ICGs detected from excitatory neuron cell pair interactions showed significant enrichment in various synapse-related GO terms, with the most significant enrichment in glutamatergic synapses. We show a list of significant GO terms and their p values adjusted for multiple testing. f. A boxplot of directional association scores of all tested genes in and out of the glutamatergic synapse gene set. g. Distance-expression profiles of the top 3 ICGs in the glutamatergic synapse gene set.

**Supplementary Table S1:** Several existing methods for cell-cell interaction analysis leveraging SRT data. We reviewed each method’s analytical framework, modeling approach, and statistical inference procedure using a set of criteria. Giotto^4^ and NCEM ^5^ are directly comparable to QuadST as they allow the identification of ICGs for a particular cell type pair’s interaction of interest. On the other hand, other methods focus either on identifying ICGs without specifying a particular cell type pair of interest^6,7^ or on identifying the patterns of cell-cell interactions using ligand and receptor pairs or a set of highly variable genes ^8–10^. Note that Giotto^4^ and NCEM ^5^ require a predetermination of distance or adjacency threshold to divide cells into a dichotomous variable of adjacent vs. non-adjacent cells, which differs from QuadST’s modeling approach. The majority of methods did not consider modeling excess zeros arising from single-cell transcriptome data and lacked robust and efficient FDR control (i.e., applicable in the presence of correlations and confounders) and covariate adjustment procedure.

|  | Giotto <sup>4</sup> | NCEM <sup>5</sup> | SVCA <sup>6</sup> | MISTy <sup>7</sup> | SpatialDM <sup>8</sup> | stLearn <sup>9</sup> | SpaOTsc <sup>10</sup> |
| --- | --- | --- | --- | --- | --- | --- | --- |
| Analytical framework | Two group comparison test | Graph neural network algorithm | Gaussian random effect model | Random forest | Descriptive spatial statistics | Spatial enrichment test | Optimal transport/Random forest |
| Detecting ICGs for a specific cell type pair's interaction | ✓ | ✓ |  |  |  |  |  |
| Dichotomizing cell-cell distance | ✓ | ✓ |  |  |  | ✓ |  |
| Lacking zero-inflation in the model | ✓ | ✓ | ✓ | ✓ | ✓ | ✓ | ✓ |
| Lacking covariate adjustment (On the fly) | ✓ |  | ✓ | ✓ | ✓ | ✓ | ✓ |
| Lacking robust and efficient FDR controlling | ✓ | ✓ | ✓ | ✓ | ✓ | ✓ | ✓ |
| Relying on known ligand-receptor pairs |  |  |  |  | ✓ | ✓ |  |

To gain insight into the biological roles of ICGs, we examined existing databases for ligands, receptors^13^, transcription factors, and cofactors^14^. We found that the detected ICGs consisted of 5.3% of ligands and receptors and 5.0% of transcription factors and cofactors in seqFISH+ and 19.5% of ligands and receptors and 8.7% of transcription factors and cofactors in MERFISH datasets (Fig. 2c). This suggests that ICGs consist of genes with various functions and cellular locations that are presumed to play a role in cell-cell interactions. We highlighted two ligand-receptor ICG pairs, including *Ctsd* -*Lrp1* and *Lrrc4* -*Ntng2*. They are detected in astrocyte-excitatory neuron/excitatory neuron-astrocyte cell pair and excitatory neuron-excitatory neuron cell pair interactions (Fig. 2b) and showed their expression-distance profiles (Fig. 2d). We note that *Lrp1* is highly expressed in neurons and glia cells and is known to function as a large multi-functional receptor that regulates the endocytosis^15^, and *Ctsd* is among the previously reported *Lrp1* ligands^16^. Additionally, the *Lrrc4* -*Ntng2* pair has been previously reported to show a receptor–ligand interaction in mouse brain in the auditory pathway ^17^.

To further understand the role of ICGs in cell-cell interactions, we tested their enrichment in GO cellular components among all anchor-neighbor cell pairs with at least 10 ICGs detected (Methods). We found that ICGs in excitatory neuron-excitatory neuron cell pairs showed a significant enrichment in synapse-related GO terms, with the most significant enrichment in glutamatergic synapses (Fig. 2e). We show a boxplot of directional association scores of all genes, grouped by membership in and out of the glutamatergic synapse gene set (Fig. 2f). Next, the expression-distance profiles of the top 3 ICGs in the glutamatergic synapse gene set are shown (Fig. 2g). The top 3 ICGs, *Cplx1, Syngap1*, and *Nsmf*, play important roles in interactions between excitatory neurons. *Cplx1* is a small synaptic protein regulating synaptic vesicle fusion ^18^, *Syngap1* encodes a Ras GTPase activating protein known for its role in modulating synaptic plasticity and neuronal homeostasis^19^, and *Nsmf* encodes the Jacob protein facilitating the transmission of signals from both synaptic and extrasynaptic NMDA receptors to the nucleus ^20^.

In addition to the detection of ICGs, QuadST provides a cell-cell interaction distance for interacting anchor-neighbor cell pairs (Supp. Fig. S1b). We found that the cell-cell interaction distance distributes approximately on the order of 10 to 100 *μm* in the seqFISH+ and MERFISH datasets. While this result is in line with the cell-cell interaction distance reported by NCEM ^5^, our analysis further indicates that cell-cell interaction distances may vary between different cell-type pairs, reflecting the presumed differences in signaling processes.

## Discussion

We introduced QuadST for the robust and powerful identification of interaction-changed genes (ICGs) for a specific cell type pair interaction using single-cell SRT data. We demonstrated that QuadST controls the FDR at a prescribed level and shows consistently comparable power or better power than other compared methods across various simulation settings. We showed that this consistent performance of QuadST is due to its statistical modeling and inference approaches that improve upon other methods. This includes i) modeling cell-cell distance as a continuous response variable instead of a dichotomous predictor variable, modeling the response variable with entire cell-cell distance quantiles instead of that with mean cell-cell distance only, and iii) testing hypotheses based on empirical FDR control procedures applicable in the presence of correlation and confounding variables. Moreover, in the application of QuadST to SRT data, we incorporated the information about excess zeros arising from the transcriptome data as another predictor in the distance-expression association model of QuadST. This needed a revision of the hypothesis testing procedure correspondingly, which can further augment the power of the QuadST and allow us to determine the direction of association for ICGs. With the application of QuadST to multiple SRT datasets, we have shown that QuadST’s unbiased inference can provide novel insights into cell-cell interactions. First, the majority of major neuronal and nonneuronal cell types are involved in cell-cell interactions. Second, ICGs consist of genes with various gene functions and cellular locations, presumed to play a role in cell-cell interactions directly and indirectly. Third, ICGs represent gene sets pertinent to a specific cell type pair’s interaction. For example, ICGs in excitatory neuron cell interactions showed significant enrichment in GO terms related to synaptic functions. Fourth, ICGs can either increase or decrease their expression levels in the presence of cell-cell interactions. In addition to the identification of ICGs involved in specific cell-type pair interactions, QuadST can offer an approximate range of cell-cell interaction distances, which may have implications for signaling processes mediating specific cell-type pair interactions. These results suggest that QuadST provides robust and powerful identification of genes involved in a specific cell type pair’s interaction and novel insights into cell-cell interaction processes from SRT data.

We expect QuadST to be an increasingly appealing approach for identifying cell-cell interactions and their contributions to complex human diseases as SRT technologies and their companion analytical tools advance. First, QuadST is well suited to handle a large number of cells and genes in high-throughput SRT data. It borrows information across genes and cells to provide FDR-controlled identifications, whose robustness and power will be enhanced rather than penalized by the growing availability of the data. Second, the improvement in cell type classification will augment QuadST’s ability to identify interactions between specific cell subpopulations. Third, with increased spatial resolution and reduced image segmentation error of SRT data, QuadST can offer a better estimation of cell-cell interaction distance. Forth, QuadST has the flexibility to define cell-cell distance considering the shape of a cell. While we determined distance by calculating cell center-to-center distance in current analysis, its analytical framework can readily be integrated with image segmentation techniques to model differently measured distances, such as surface-to-surface distance. Finally, with the emerging subcellular resolution data, QuadST’s approach can be extended to identify the subcellular localization of ICGs. For example, QuadST can model and test the association between cell-cell distance and the presence within a particular subcellular location of ICGs. In addition, we can consider other possible methodologic extensions toward a deeper understanding of cell-cell interaction processes (e.g., identification of regulatory processes among ICGs).

Since cell-cell interactions can affect the function of tissues and organs, we expect that QuadST has potential applications in the identification of specific cell types and genes that play a role in cell-cell interactions under normal and abnormal functions in various tissues and organs. For instance, it is known that several genetic loci associated with the risk of late-onset Alzheimer’s disease (AD) involve genes exclusively or highly expressed in microglia ^21,22^, which play a protective role in the function of neurons in various ways ^23–26^ in healthy brains. However, genes and pathways that regulate microglia-neuron interactions are poorly understood in healthy brains and in AD. Therefore, we can apply QuadST as a tool to investigate genes and pathways involved in microglia-neuron interactions in healthy brains and AD as well as to investigate genes and pathways involved in specific cell-cell interactions relevant for healthy and diseased states of many other tissues and organs^27,28^.

**Supplementary Figure S1:**
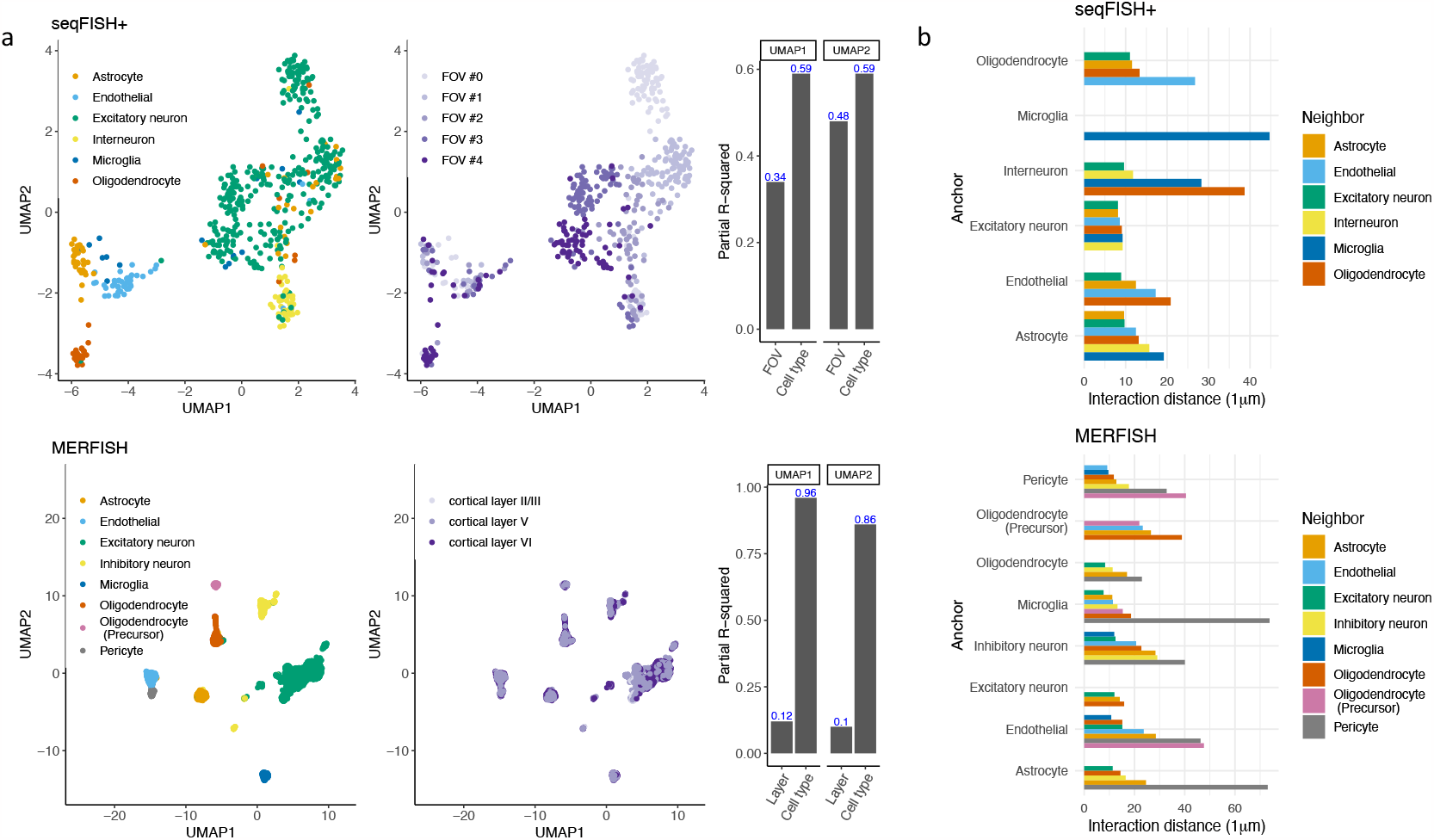
The UMAP plot and cell-cell interaction distance of two SRT datasets. a. The UMAP plot of seqFISH+ and MERFISH datasets. Cells cluster primarily by cell type in both datasets (first column) and secondarily by FOV and cortical layers (second column) in seqFISH+ and MERFISH datasets respectively, which were adjusted for in our analysis. The partial R-squared of each covariate for UMAP coordinates is shown along with UMAP figures (third column). b. Cell-cell interaction distances for interacting anchor-neighbor cell pairs. The cell-cell interaction distance was approximately on the order of 10 to 100 *μm* in the seqFISH+ and MERFISH datasets.

## Methods

### QuadST

#### Overview

The statistical framework of QuadST consists of three key steps: (1) Create an anchor-neighbor integrated matrix. (2) Model anchor-neighbor cell–cell distance quantile with the gene expression level of anchor cells. (3) Identify ICGs for a given anchor-neighbor cell pair based on a hypothesis testing with empirical distribution applicable in the presence of correlations and confounding variables (Fig. 1a).

#### Anchor-neighbor integrated matrix

Suppose a SRT data contains expression levels of *G* genes in *N* cells. These cells can be categorized into *K* cell types based on their expression (and sometimes spatial) profiles ^4^. For cells in each cell type *k ∈* (1, …, *K*), we are interested in identifying ICGs in the presence of cell-cell interaction with the same or other cell type *l∈* (1, …, *K*). To this end, we define anchor-neighbor integrated matrix for each of all *K* by *K* anchor-neighbor cell type pairs. Specifically, we define 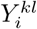 as the distance from cell *i ∈* (1, …, *N*_*k*_) in cell type *k* to its most proximal neighbor cell in cell type *l*, where *N*_*k*_ is the number of cells for cell type *k*. We also define 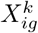 as the expression level of gene *g ∈* (1, …*G*) and 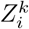 as as the cell-level covariates for each cell *i* in cell type *k* respectively. For simplicity, we will use *Y*_*i*_, *X*_*ig*_, and *Z*_*i*_ in the following section, omitting anchor-neighbor cell type pair index (*kl*) as our analysis applies equivalently to each of *K* by *K* cell type pairs.

#### Distance-quantile based model

Our statistical model is built based on the following two assumptions: First, anchor cells in close proximity to neighbor cells are more likely to be involved in cell-cell interaction. Second, anchor cells’ gene expression level can change due to cell-cell interaction. Under these assumptions, we can reason that in the presence of cell-cell interaction, anchor cells proximal to neighbor cells can show a change in gene expression level, the effect of which can be pronounced within and in the vicinity of a certain cell-cell interaction distance. Therefore, to identify ICGs, we can model and test association between conditional cell-cell distance quantile and gene expression level. Modelling multiple quantile levels across the entire cell-cell distance distribution allows us to assess the association without the need of knowing a specific quantile level where cell-cell interaction occurs. We can then write the model for a gene *g* at a quantile level *τ ∈* (0, 1) as follows.

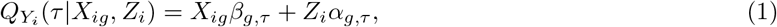

where *β*_*g,τ*_ and *α*_*g,τ*_ capture the effects of gene expression level and covariates of gene *g* at a given quantile level *τ*, respectively.

It is well known that single cell transcriptome data contains excessive zeros arising from biological and technical sources. It remains an open question as to how to properly incorporate them in a statistical model^29^. In this work, we consider that the observed number of nonzeros provides a complementary information about the level of gene expression and thus, expand our model to include nonzero indicator as an additional predictor.

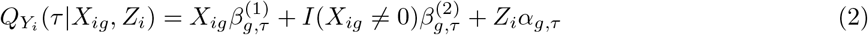

where *I*(.) is an indicator function of whether gene expression level is nonzero. 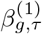and 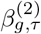are parameters capturing the effects of gene expression level and nonzeros of gene g at a given quantile level *τ*, respectively.

For both of equations, (1) and (2), we calculated the test statistic to test whether the association parameter *β*_*g,τ*_ = 0 or the combined association of two parameters, 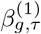and 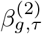 is jointly equal to zero, considering their directions of association. For the former, we used QRank method^30^ and for the latter, we used Signed-QRank as described in *Details on signed multivariable QRank (Signed-QRank)*.

#### Identification of ICGs

To infer cell-cell interaction and its ICGs, we first perform quantile regression at a series of evenly spaced cell-cell distance quantiles ***τ*** = (*τ*_1_, *τ*_2_, …, *τ*_*J*_) for each gene *g ∈* (1, 2, …, *G*). This yields a *G× J* p-value matrix, with each element 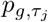 capturing association at each quantile level *τ*_*j*_ for each gene *g*. We then combine gene-specific p-values across *j* highest and lowest quantiles symmetrically around the median (*τ* = 0.5). The combined p-values are denoted as 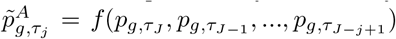 for the upper quantiles and 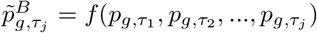for the lower quantiles, respectively, with *f* (.) referring to Cauchy combination test^31^.

Since we assume that cell-cell interaction is stronger for cells in close proximity compared to those further apart, we compare 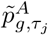and 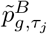to identify ICGs. Specifically, at a specific quantile level *τ*_*j*_, we calculate *T*_*j*_(*c*) as the ratio of genes with p-values below a small positive cutoff *c* for the upper and lower tails, as such

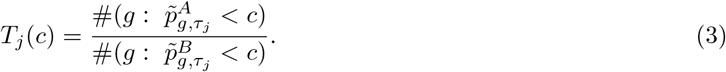

A ratio close to 1 indicates a similar number of genes with small p-values for both close and distant cells, suggesting a lack of cell-cell interactions. Conversely, a small ratio suggests more genes with small p-values in nearby cells than distant cells, indicating the presence of cell-cell interactions. Therefore, an evaluation *T*_*j*_(*c*) can be used to infer the presence of cell-cell interaction and its ICGs.

It is critical to note that *T*_*j*_(*c*) provides an upper bound of the false discovery rate (FDR) under two simple conditions:

1. The null distributions of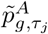 and 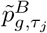are symmetric, such that 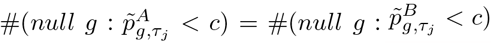.
2. The alternative distribution is more extreme than the null distribution, resulting in 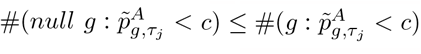.

In this context, the empirically defined FDR (eFDR^32^) can be expressed as:

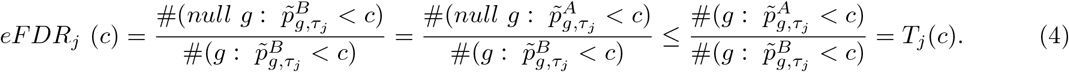

This means that *T*_*j*_(*c*) has a highly attractive feature that it can provide robust and efficient FDR control for ICG identification, even in the presence of complex p-value behaviors in real data. First, *T*_*j*_(*c*) does not assume well-behaved distributions of 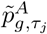and 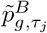 under the null. In scenarios involving unmeasured confounders, the distance-quantile based models might generate small p-values under the null. However, by comparing the relative values of 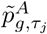and 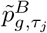 instead of comparing each of them to a theoretical distribution, *T*_*j*_(*c*) ensures FDR control without relying on specific assumptions about the p-value distributions. Second, *T*_*j*_(*c*) does not assume any dependency structures for 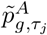and 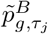. This makes it robust under unknown gene-gene and quantile-quantile correlations. Unlike many existing FDR control procedures that rely on assuming known distributions or correlation structures, *T*_*j*_(*c*) provides flexibility and robustness in handling complex and unknown p-value behaviors.

To identify ICGs at a desired nominal eFDR level *α*, we search for the p-value threshold *c* as

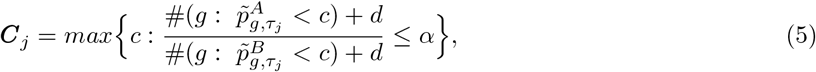

where *d >* 0 is a small constant to make eFDR control conservative.

Finding the appropriate threshold ***C***_*j*_ indicates that cell-cell interaction exists at or below the desired nominal FDR level *α* at the quantile level *τ*_*j*_. Then, the significant genes can be determined as

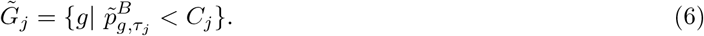

ICGs are the significant genes at quantile level *I*, i.e. 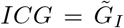, where *I* maximizes the number of identified genes, i.e. 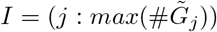. Meanwhile, the cell-cell interaction distance is the distance corresponding to quantile level *I*.

#### Details on signed multivariable QRank (Signed-QRank)

In joint testing of multiple parameters, rank-score test statistics do not consider the directions of association of each parameter. However, in our application, the direction consistency of *β*^(1)^*g, τ* and *β*^(2)^*g, τ* is crucial for characterizing the expression-distance associations. To address this, we propose an extension of our previous method QRank, called Signed-QRank, that allows for considering the directionality of the signals in joint testing of multiple parameters.

Specifically, we let *Y* denote the outcome vector with length *n, X* denote the *n* × *p* matrix for multiple primary predictors, and *Z* denote the *n × q* matrix for covariates including intercept (design matrix under the null). Following the reference ^33^, we first project *X* to the column space of *Z* to obtain orthogonal signals of covariates, such that *X*^*\**^ = *X* − *Z*(*Z*^*T*^ *Z*)^−1^*Z*^*T*^ *X*. Then, the quantile rank score function can be defined as 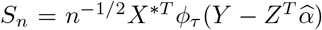, where *ϕ*_*τ*_ (*u*) = *τ* − *I*(*u <* 0) is an asymmetric sign function, and 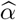 is the estimated coefficient under the null. Under the null hypothesis, *S*_*n*_ is a *p ×* 1 matrix following a normal distribution with mean zero and variance 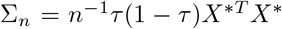. In Signed-QRank, we standardize *S*_*n*_ and take an average across the *p* predictors, such that 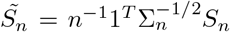. The 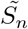 is our test statistics that follows *N* (0, 1) under the null.

### Simulation studies

We designed simulation studies to evaluate the performance of QuadST in comparison with other methods. The simulation studies allow us to estimate the power and FDR given a ground truth set of interacting cells and IGCs in three settings.

#### Overall Design

We simulated SRT data under three different settings that represented different spatial organization of cells and their expression levels. Across all settings, we simulated a total of 5000 cells equally divided by 5 different cell types. From these 5 cell types, one cell type was chosen as the anchor cell, and another cell type was selected as the neighbor cell. In each of anchor cell, we simulated 4000 genes’ expression levels. These genes were block-wise correlated, with a block size of 100. Within each block, gene expression levels followed a multivariate normal distribution with an autocorrelation structure and with homogenous variance. The autocorrelation coefficient between two adjacent genes was 0.7. We also assumed that 10% of the anchor cells, which were most proximal to paired neighbor cells, were involved in two cell type pair’s interactions. We perturbed the expression levels of 10% genes (i.e., 4 blocks out of 40 blocks) to simulate the cell-cell interaction between anchor and neighbor cells.

#### Simulated Settings

The following describes each simulation setting:

#### Setting 1: Spatially Uniform

This setting was designed to simulate a SRT data without a particular spatial pattern in both cell composition and gene expression level. To this end, both anchor and neighbor cells were uniformly distributed in a unit square. Under the null of no cell-cell interaction, the mean of gene expression level was set at 0 with its variance at 1. Under the alternative, the details of simulated ICGs were described in the next section.

#### Setting 2: Spatially Variable Cells and Genes

This setting was designed to simulate a SRT data with spatial variation in both cell composition and gene expression level. Specifically, we considered a circumstance where anchor cells’ gene expression levels could systematically increase or decrease with distance from neighbor cells without the presence of cell-cell interaction. To this end, we created 4 spatial regions dividing a unit square into 4 equal-sized squares and denote them as *R*_1_, *R*_2_, *R*_3_, and *R*_4_ respectively. We then modified gene expression levels and cell-type composition depending on spatial regions: i) We increased the mean of anchor cell’s gene expression level in a spatial region *R*_4_ under the null by 0.2, while the mean of anchor cell’s gene expression level in all other regions was kept at 0. ii) We restricted a paired neighbor cells’ presence only within spatial regions, *R*_1_, *R*_2_, *andR*_3_. This restriction increased anchor-neighbor cell distance for anchor cells in *R*_4_ compared to those in *R*_1_, *R*_2_, *andR*_3_. With these modifications, anchor cells located in a particular spatial region (*R*_4_) could have elevated gene expression level and farther cell-cell distance from neighbor cells without the presence of cell-cell interaction, which made spatial region serve as a confounding factor of the expression-distance association.

#### Setting 3: Spatially Variable Cells only

This setting was designed to simulate a SRT data with spatial variation only in cell composition. In this setting, we simulated the spatial distribution of cells in 4 spatial regions like Setting 2. However, we kept the mean expression levels of anchor cells the same for all spatial regions (*R*_1_, *R*_2_, *R*_3_, and *R*_4_) under the null. Thus, anchor cells located in a particular spatial region (*R*_4_) had the same expression levels with other spatial regions (*R*_1_, *R*_2_, and *R*_3_). Therefore, the spatial variation in cell composition alone did not serve as a confounding factor.

#### Simulated ICGs

The ICGs were given by a few different distance-expression association patterns characterized by how the mean and variance of gene expression level vary with cell–cell distance. Specifically, we simulated four different distance-expression association patterns in four different blocks as follows:

#### Block 1: Constant Mean Shift

This pattern represents an “on/off” interaction, where cell-cell interaction is turned “off” above a certain cell-cell distance threshold and “on” below the threshold. Under the interaction, the effect is a constant mean shift with parameter *μ*_*δ*_, where *μ*_*δ*_ ∼ *N* (0.4, 0.2).

#### Block 2: Distance-dependent Mean Shift

This pattern represents a “dosage” interaction, where cell-cell interaction increases as cells become closer after being turned “on” at a distance threshold. The mean shift is generated as a piece linear function of distance (*Y*), such that *μ*_*δ*_(*Y*) = *a*(*Y* − *d*_*c*_)*I*(*Y < d*_*c*_), where *d*_*c*_ is the distance cutoff for 10% of cells and *a* ∼ *N* (−150, 75) is the slope for dosage effect.

#### Block 3: Constant Mean Shift with Increased Variance

Qualitative Mean/Scale Change. Similar to the “on/off” pattern in Block 1, but interaction not only impacts mean, but also the variance of gene expression. We simulated the variance of interacting genes *var*_*δ*_ = 2.

#### Block 4: Distance-dependent Mean Shift with Increased Variance

Similar to the “dosage” pattern in Block 2, but interaction not only impacts mean, but also the variance of gene expression. We simulated the variance of interacting genes *var*_*δ*_ = 2.

#### Compared methods

We compared QuadST with the three other methods, including Giotto^4^ and NCEM ^5^ from existing methods (Supplementary Table S1) and “Linear regression” derived by us.

#### Giotto

Giotto’s ICG analysis required the adjacency between two cells from anchor-neighbor cell pair, either determined by a certain distance threshold or network analysis. Two different approaches, Giotto *A* and Giotto *B*, were considered in determining the adjacency. Giotto *A* matched the distance threshold with the simulated ground truth cell-cell interaction distance. Giotto *B* used a distance threshold that was 1.4 times of the simulated ground truth cell-cell interaction distance. This increase in the distance threshold approximately doubled the adjacent area in which anchor-neighbor cell pairs were preassigned as putative interactors. ICGs were identified by t-test between neighbors and non-neighbors and p-values were adjusted for multiple testing (i.e., FDR *<* 0.1).

#### NCEM

NCEM was built on a spatial neural network of cells. The adjacency between two cells from anchor-neighbor cell pair can be determined by a certain distance threshold. Like Giotto, two different approaches, Linear NCEM *A* and Linear NCEM *B*, were tested in determining the adjacency. Linear NCEM *A* matched the distance threshold with the simulated ground truth cell-cell interaction distance. Linear NCEM *B* used a distance threshold that was 1.4 times of the simulated ground truth cell-cell interaction distance. Although NCEM can search cell-cell interaction distance by maximizing the model evaluation criteria (i.e., *R*^2^ between predicted and observed gene expression levels), it is possible that the identified cell-cell interaction distance differs from the ground truth interaction distance. We used a custom code to implement a Linear NCEM spatial model for an anchor-neighbor cell pair. The model equation can be expressed as follows: *E*[*Y*_*ig*_] = *X*_*i*_*β*_0,*g*_ +*S*_*i*_*β*_1,*g*_ +*Z*_*i*_*α*_*g*_, where *Y*_*ig*_ : the gene expression level of a gene *g ∈* (1, …, *G*) in an anchor cell *i ∈* (1, …, *N*), *X*_*i*_ : the design matrix indicating a cell type of cell *i, S*_*i*_ : the design matrix indicating the presence of neighbor cells of cell i within a distance threshold, and *Z*_*i*_ : the design matrix for cell’s covariates. The *β*_0,*g*_ and *β*_1,*g*_ captures the effect of cell type and interaction with neighbor cells while *α*_*g*_ capture the covariate effect. ICGs were identified by the significance of coefficient *β*_1,*g*_ for each gene *g* and p-values were adjusted for multiple testing (i.e., FDR *<* 0.1).

#### Linear regression

Like QuadST, this method models cell–cell distance as a response and gene expression level as a predictor. The model can be expressed *E*[*Y*_*i*_] = *X*_*ig*_*β*_*g*_ + *Z*_*i*_*α*_*g*_, where *Y*_*i*_ is the distance of an anchor cell *i ∈* (1, …, *N*) from the most proximal neighbor cell, *X*_*ig*_ : the expression level of a gene *gin*(1, …, *G*) in a cell *i*, and *Z*_*i*_ is the cell level covariate of cell *i*. The *β*_*g*_ and *α*_*g*_ capture the effects of gene expression level and covariates of gene *g*. ICGs were identified by the significance of coefficient *β*_*g*_ for each gene *g* and p-values were adjusted for multiple testing (i.e., FDR *<* 0.1).

### Data analysis

#### seqFISH+ dataset

High-throughput single-cell SRT data obtained from the cortex of a 23-day-old male mouse (C57BL/6J) using seqFISH+ were downloaded for analysis^11^. It included spatial map and expression levels for 10,000 genes in 511 cells, imaged from 5 field of views (FOVs). The cells were categorized by the original study into six major cell types ^11^: excitatory neuron (n=325), interneuron (n=42), astrocyte (n=54), endothelial (n=45), oligodendrocyte (n=29), and microglia (n=16).

#### MERFISH dataset

We also analyzed the single-cell SRT data from mouse frontal cortex, profiled using MERFISH^12^. The analysis focused on a 4 week-old female mouse (C57BL/6J) similar in age to seqFISH+ data. The data consists of gene expression levels for 374 genes in 13,745 cells collected from cortical layers II-V. These cells have been categorized into eight major cell types: excitatory neuron (n= 7129), inhibitory neuron (n=1161), astrocyte (n=1333), microglia (n=787), oligodendrocyte (n=1153), oligodendrocyte-precursor (n=462), endothelial (n=1360), and pericyte (n=360) 12. We used each cell’s spatial coordinates as provided in CELL x GENE repository (shown in the key resource table of the reference^12^).

#### Dataset preprocessing

To preprocess seqFISH+ data, we stitched cell’s coordinates obtained from 5 FOVs to provide global cell spatial coordinates ^11^. We normalized gene counts across all cells using scran ^34^, which adjusted for cell-specific bias due to cell-to-cell difference in library size and capture efficiency. Following the practice in the source ^11,35^, we kept for analysis highly-expressed genes whose expression levels belong to top 25% quantile in each cell type. To preprocess MERFISH data, we normalized gene counts using scran ^34^ to adjust for cell-specific bias and kept genes whose expression levels belong to top 75% quantile in each cell type. For each dataset, the cell specific-bias adjusted counts were transformed to a normal distribution as follows: i) we first calculated the empirical cumulative distribution function (ecdf R function) of the count data to transform the counts to values between 0 and 1. ii) we used inverse cumulative distribution function of a normal distribution to transform values between 0 and 1 to a normal distribution (qnorm R function). iii)we shifted the minimum of a normal distribution to 0.

#### UMAP and covariate effect analysis

We performed UMAP analysis on gene expression data and examined the effect of covariates on the gene expression profiles of cells from each SRT dataset. To this end, we calculate partial *R*^2^ of each covariate with each UMAP coordinate using rsq R package, in a generalized linear model where each UMAP coordinate of each cell is considered as a response and covariates of interest as predictors in each SRT dataset. We used umap R package to obtain UMAP coordinates of cells.

#### Running QuadST for SRT dataset

We ran QuadST on each of the two preprocessed SRT datasets. First, we created anchor-neighbor integrated matrices for all pairs of the same and different cell types. Second, we used the distance-quantile based model given by Eq. (2) to test distance-expression association at a series of evenly spaced distance quantiles. The number of quantile levels was chosen adaptively to ensure that there were at least 5 samples between two selected quantile levels with the maximum number of quantile levels set at 49. Third, we identified ICGs based for each cell-type pair with 10% empirical FDR control procedure.

#### Calculating directional association score

In order to determine the direction of association, i.e., whether gene expression increases or decreases around cell-cell interaction distance, we first calculated the test statistics associated with two primary parameters and combined them into a single z-score considering their correlation as detailed in Signed-QRank section. The sign of z-score indicate whether gene expression increases or decrease with cell-cell distance. To combine the level of association and the direction, we calculated the directional association scores as -log10 (“adjusted Cauchy combination test p-value”)sign(“z-score”) at the interaction quantile level (I) of each anchor-neighbor cell type pairs.

#### GO cellular component enrichment analysis

To test GO cellular component terms enriched for ICGs, we performed Wilcoxon Rank Sum test comparing the ranks of p-values of genes overlapping vs. nonoverlapping with a given GO cellular component gene set for all cell type pairs with at least 10 ICGs detected. We downloaded GO cellular component gene sets for mouse (m5.go.cc.v2023.1.Mm.symbols.gmt) from GSEA website (https://www.gsea-msigdb.org/gsea/msigdb/mouse/collections.jsp). To identify significantly enriched GO cellular component terms, we adjusted Wilcoxon Rank Sum test p-values for multiple testing (i.e., FDR *<* 0.1).

#### ICG function annotation using existing databases

To examine known biological roles of ICGs, we looked up in existing databases on ligands and receptors^13^ as well as transcription factors and cofactors^14^. We obtained a set of mouse ligands and receptors (mouse lr pair.txt) from CellTalkDB website (http://tcm.zju.edu.cn/celltalkdb/) and mouse transcription factors and cofactors from a supplementary table (41467 2017 BFncomms15089 MOESM3449 ESM.xlsx) of the reference ^14^.

## Acknowledgements

This work was supported by NIH grants: R03AG075567 and RF1MH133703.

